# Genome-wide survey of the F-box/Kelch (FBK) members and molecular identification of a novel FBK gene *TaAFR* in wheat

**DOI:** 10.1101/2021.04.08.438931

**Authors:** Chunru Wei, Weiquan Zhao, Runqiao Fan, Yuyu Meng, Yiming Yang, Xiaodong Wang, Nora A. Foroud, Daqun Liu, Xiumei Yu

**Affiliations:** College of Life Sciences/Key Laboratory of Hebei Province for Plant Physiology and Molecular Pathology, Hebei Agricultural University, Baoding, Hebei, China; Technological Innovation Centre for Biological Control of Crop Diseases and Insect Pests of Hebei Province, Hebei Agricultural University, Baoding, Hebei, China; Lethbridge Research and Development Centre, Agriculture and Agri-Food Canada, Lethbridge, Alberta, Canada

**Keywords:** Wheat F-box/Kelch, Genome-wide survey, AFR, Expression pattern, Protein interaction

## Abstract

F-box proteins play critical roles in plant responses to biotic/abiotic stresses. In the present study, a total of 68 wheat F-box/Kelch (*TaFBK*) gene sequences encoding for 74 proteins were obtained in a genome-wide survey against EnsemblPlants. The 74 TaFBK proteins were divided into 5 categories based on their domain structures. The FBK proteins from wheat, Arabidopsis, and three other cereal species were grouped into 7 clades, and the number of Kelch domains present was their key clustering criterion. Sixty-eight *TaFBK* genes were unevenly distributed on 21 chromosomes. Most of TaFBKs were predicted to localize in the nucleus and cytoplasm. *In silico* analysis of a digital PCR revealed that *TaFBKs* were expressed at multiple developmental stages and tissues, and in response to drought and/or heat stresses. The *TaFBK19* gene, a homologous to the *Attenuated Far-Red Response* (*AFR*) genes in other plant species, and hence named *TaAFR*, was selected for further analysis. The gene was isolated from the wheat line TcLr15 and its expression evaluated by quantitative real-time PCR. *TaAFR* transcripts were primarily detected in wheat leaves, and its expression was found to be regulated by various abiotic and biotic stresses as well as plant signaling hormones. Of particular interest, *TaAFR* expression was differentially regulated in the compatible *vs* incompatible wheat leaf rust reaction. Subcellular localization studies showed that TaAFR accumulates in the nucleus and cytoplasm. Three TaAFR-interacting proteins were identified experimentally: Skp1/ASK1-like protein (Skp1), ADP-ribosylation factor 2-like isoform X1 (ARL2) and phenylalanine ammonia-lyase (PAL). Further analysis revealed that the Skp1 protein interacted specifically with the F-box domain of TaAFR, while ARL2 and PAL were recognized by the Kelch domain. The data presented herein provides a solid foundation from which the function and metabolic network of TaAFR and other wheat FBKs can be further explored.

## Introduction

In eukaryotes, the ubiquitin/26S proteasome system (UPS) is responsible for the selective degradation of most intracellular proteins [1]. Together with Suppressor of Kinetochore Protein 1 (SKP1), Cullin 1 (CUL1) and Ring-Box 1 (RBX1), F-box proteins form a ubiquitin ligase complex, where it plays the critical role of recruiting substrates to the UPS [2]. F-box proteins carry one or more 40-50 residue F-box/F-box-like domains in their N-terminus that are in charge of binding to Skp1/Skp1-like proteins [3]. Meanwhile, one or more additional conserved domains involved in substrate specificity can be found downstream of the F-box/F-box-like domain(s), such as Kelch repeats, Leucine Rich Repeat (LRR) and WD40-repeats [4]. The F-box members form a large family of proteins in plants. Within the F-box family, the Kelch subfamily is one of the large groups and F-box/Kelch (FBK) proteins are almost exclusively found in plants. The Kelch domain, originally identified in *Drosophila* mutants consists of 44-56 residues [5], and one or more Kelch domains can be found in an FBK protein.

The size of the FBK subfamily varies depending on the plant species. In 2009, Xu et al. reported the identification of 96 FBKs in Arabidopsis, along with 27 and 35 FBKs from rice and poplar, respectively [6]. Using this information, Schumann et al. went on to identify additional FBKs in numerous species, and found 103, 68 and 36 FBKs from the dicot species, *Arabidopsis thaliana*, *Populus trichocarpa* and *Vitis vinifera*, respectively, 44 and 39 FBKs in the monocot species, *Sorghum bicolor* and *Oryza sativa*, respectively, and 71 and 46 FBKs in the non-seed embryophytes *Physcomitrella patens* and *Selaginella moellendorffii*, respectively [7]. The former study reported that the FBK subfamily altered their protein structures by increasing or decreasing the number of exons, and the subfamily size was expanded primarily *via* tandem duplications [6]. The FBKs have been found to participate in biological clock regulation, photomorphogenesis, phenylpropanoid and pigmentation biosynthesis and biotic stress responses [6, 8–12]. While the FBKs subfamily exists in plants in relatively high numbers, and participates in many important biological processes, no systematic studies of the FBK subfamily have previously been reported in hexaploid wheat species.

To initiate FBK research in hexaploid wheat and to further our understanding of their role in various biological processes, a genome-wide identification study of this subfamily of F-box proteins and a systemic analysis of protein structure, phylogenetic relationship, chromosome distribution, and expression patterns in response to different stresses are presented herein. Sixty-eight genes encoding 74 wheat FBK (TaFBK) proteins were identified, and an analysis of protein structure, phylogenetic relationship, chromosome distribution, and expression patterns in response to different stimuli and stresses is presented. *In silico* expression analysis revealed that these genes were differentially regulated in response to drought and heat stress. One gene, *TaFBK19*, which showed similarities to the *Attenuated Far-Red Response* (*AFR*) gene, was selected for further investigations, and is described here as *TaAFR*.

AFR F-box genes are involved in light signaling but have also been shown to participate in plant stress responses. Through the course of its cultivation, wheat is subjected to many kinds of environmental and biotic stresses including salt, drought, cold, heavy metals and various pathogens. These stresses can affect crop productivity and yield, which can be mitigated if a timely and appropriate stress response is mounted in the plant. To determine whether *TaAFR* is involved in the plant’s response to different stress stimuli, the wheat line TcLr15 was exposed to leaf rust pathogen, salt, drought and H_2_O_2_, salicylic acid (SA), abscisic acid (ABA) and methyl-jasmonate (MeJA), and changes in gene expression were assessed by quantitative real-time PCR (qRT-PCR). Subcellular localization of TaAFR was experimentally determined, and it’s interactions with other proteins was investigated using a combination of yeast-2-hybrid (Y2H), bimolecular fluorescence complementation (BiFC) and co-immunoprecipitation (Co-IP) assays. While providing a glimpse into the function of TaAFR and other FBKs in wheat, the results presented herein build the foundation to further dissect the function and metabolic network of this important gene family.

## Materials and Methods

### Genome-wide survey of wheat FBKs

#### Database search, sequence analysis and classification of wheat FBKs

The Hidden Markov Model (HMM) profiles of the F-box domain (PF00646, PF15966), F-box-like domain (PF12937, PF13013) and Kelch domain (PF01344, PF07646, PF13415, PF13418, PF13854, PF13964) were obtained from Pfam (http://pfam.xfam.org/). To identify wheat FBKs, the HMMER3.1b2 software was first used to search for F-box and F-box-like domains encoded in wheat genes deposited in the IWGSC (Wheat Genome Sequencing Consortium) RefSeq v1.0 wheat database downloaded from EnsemblPlants (https://plants.ensembl.org/index.html) (E value cut-off of 1.0) [13], and TBtools (v0.6673) was used to extract the target sequences. Sequences encoding F-box and F-box-like domains were further screened for the presence of one or more Kelch domains (E value cut-off of 1.0). Finally, Pfam, SMART (http://smart.-heidelberg.de/) and HMMER (web version 2.25.0, https://www.ebi.ac.uk/Tools/hmmer/) were adopted to confirm the presence of both the F-box (or F-box-like) and Kelch domains in each FBK protein identified, E value <1.0.; sequences that did not meet this criterion were removed.

The predicted isoelectric point (*p*I) and molecular weight (MW) of the putative wheat FBKs (TaFBKs) were computed at ExPASy (https://web.expasy.org/compute_pi/). The intron-exon organization of wheat FBKs was obtained from EnsemblPlants. Subcellular localizations were predicted using cropPAL2020 dataset (https://crop-pal.org/).

#### Analysis of conserved residues within the F-box and Kelch domains of wheat FBK proteins

The ClustalX2.0 multiple sequence alignment tool was used to align the F-box or Kelch domains extracted from the TaFBK protein sequences, and WebLogos (http://weblogo.berkeley.edu/) were generated for each of the two domains.

#### Phylogenetic analysis

In order to study the phylogenetic relationship and evolution of wheat FBKs, the obtained TaFBK sequences were compared with the orthologues of model dicot species Arabidopsis (AtFBK), and three important monocots rice (OsFBK), sorghum (SbFBK) and maize (ZmFBK). The AtFBK, OsFBK, SbFBK sequences reported by Schumann et al. and ZmFBKs reported by Jia et al. were downloaded and screened for the presence of the F-box and Kelch domains [7, 14]. Sequences that did not carry both F-box and Kelch domain(s) were removed, 94, 31, 34 and 32 FBK protein sequences were left for Arabidopsis, rice, sorghum and maize, respectively. These FBKs from wheat, Arabidopsis, rice, sorghum and maize were subsequently aligned with the ClustalX 2.0 algorithm and a phylogenetic tree was constructed by the Maximum Likelihood (ML) in MEGA7 using default parameters, with bootstrap value set to 1000 repetitions.

#### Chromosomal distribution and gene duplication analysis

The chromosomal distribution of wheat *FBK* genes were obtained from the EnsemblPlants (IWGSC RefSeq v1.0). MapDraw was used to visualize the detailed location of each *TaFBK* gene on the wheat chromosome [15]. Greater than 70% sequence similarity was set as the criterion for determining gene duplication [16]. When the maximum distance between duplicated genes on the same chromosome was smaller than 50 kb, tandem duplication and duplicated genes on different chromosomes were delimited as segmental duplication [17].

#### *In silico* expression analysis of *TaFBK* genes

*FBK* gene sequences obtained from the EnsemblPlants were input into the WheatExp wheat database (https://wheat.pw.usda.gov/WheatExp/) and searched for the corresponding gene ID. According to transcriptomics data from digital PCR experiments deposited in WheatExp, FPKM (Fragments per kilobase per million mapped reads) values of *FBKs* were obtained from 5 tissues (CS) at different development stages: leaves (z10, z23, z71), roots (z10, z13, z39), stems (z30, z32, z65), spikes (z32, z39, z65) and grains (z71, z75, z85) [18]. In order to determine the relationship between *FBKs* expression and the abiotic stress response in wheat, the FPKM values were downloaded from hexaploid bread wheat (cultivar TAM 107) treated with drought (DS), heat (HS) and drought+heat (HD) stresses [19]. TBtools (v0.6673) was used to draw a heat map according to their corresponding FPKM values.

### Molecular identification and expression patterns of *TaAFR*

The wheat FBK gene, *TaFBK19*, was selected for further analysis. This gene is similar to the Kelch containing F-box *AFR* genes from other species and is therefore described here as *TaAFR*.

#### Plant material, fungal strains and inoculum preparation

A near-isogenic wheat line of Thatcher for leaf rust resistance, TcLr15, and leaf rust strains 05-5-137③ and 05-19-43② were used in the present study. Unless otherwise specified, plants were grown in a greenhouse as described in Yu et al. [20]. Urediniospore and inoculum preparation of leaf rust pathogens were carried out as previously described [20].

#### *TaAFR* cloning

Total RNA extraction and first strand cDNA synthesis were performed as previously described [20]. A pair of gene specific primers *TaAFR*-F and *TaAFR*-R (S1 Table) and Tks Gflex™ DNA Polymerase (TaKaRa, Japan) were used to amplify the full-length coding sequences CDS amplified with Tks Gflex™ DNA Polymerase (TaKaRa, Japan) according to manufacturer’s directions with an annealing temperature of 56.4°C. The purity of the amplicon was verified by 1.2 % agarose gel electrophoresis and the product was sequenced to confirm the identity of the clone.

The TaAFR sequence was used to pull out related sequences from the NCBI transcript database using the BLASTp tool, and sequences with an expect threshold of <0.05 were aligned together with TaAFR in MEGA 7.0 and a phylogenetic tree was constructed, as described in the section on phylogenetic analysis. The TaAFR protein sequence was also analyzed using various bioinformatics tools to predict presence of signal peptides (SignalP-4.1, www.cbs.dtu.dk/services/SignalP/), transmembrane domains (TMHMM Server v. 2.0, www.cbs.dtu.dk/services/TMHMM/), and subcellular localization (cropPAL2020 dataset). The 3D structure was predicted in Phyre2 (www.sbg.bio.ic.ac.uk/phyre2/).

#### Wheat treatments and sampling for quantitative real-time PCR

Sampling of wheat for gene expression analysis was carried out in different tissues (for tissue-specific analysis) and in response to three different types of abiotic stresses and three hormone treatments, as described below. For each experiment, samples were collected from three replicates, unless otherwise specified, 3-5 samples were harvested for each replicate. Samples were flash frozen in liquid nitrogen and stored at −80 °C prior to RNA extraction.

To detect tissue-specific expression levels of the *TaAFR* gene, samples were collected from TcLr15 7-day old seedlings and adult plants grown in a pot with nutrient soil (Hebei Fengyuan, China) in a greenhouse (22°C, 16 h light/8 h dark). Roots, stems and leaves were collected at z11; pistils, stamens and flag leaves were collected at z51. A large number of pistils and stamens (50-100 mg) were sampled from wheat florets.

To assess the effect of leaf rust pathogen on *TaAFR* expression, TcLr15 plants were inoculated with 05-5-137③, 05-19-43②, or water (negative control), as previously described [20]. The inoculated leaves were harvested at 0, 6, 12, 24, 48 and 96 hours post inoculation (hpi).

The effect of abiotic stress treatments on *TaAFR* expression was evaluated in TcLr15 plants grown in Hoagland’s solution [21]. Once plants reached the three-leaf stage (z13), the Hoagland’s solution was amended with NaCl, PEG 6000 and H_2_O_2_, to a final concentration of 300 mM, 10 % and 7 mM, respectively [22–24]. The second leaves were sampled at 0, 0.5, 2, 6, 12, 24 and 48 h post-treatment. Samples were also collected at the same time points from untreated negative control plants in Hoagland’s solution.

The plant hormones, SA, ABA and MeJA, are known to be involved in both abiotic and biotic stress responses [25, 26]. To investigate the effects of 3 hormones on the expression of *TaAFR*, exogenous treatments of SA (2 mM), ABA (100 μM) and MeJA (100 μM), each dissolved in 0.1% absolute ethanol [22, 26], were applied to TcLr15 seedlings (z11) grown in pot with nutrient soil in greenhouse. The negative control plants were sprayed with 0.1% absolute ethanol. The primary leaf of each plant was sampled at 0, 0.5, 2, 6, 12, 24 and 48 h post-treatment.

#### Quantitative real-time PCR (qRT-PCR)

Total RNA was extracted from the TcLr15 samples collected in the previous section for gene expression analysis, using Biozol reagent (BioFlux, Japan) according to manufacturer’s instructions. To eliminate gDNA contamination, 2 ug of each RNA sample was treated with 1 uL gDNA Remover (TransGen, China). cDNA synthesis was carried out as described by Yu et al. [20]. qRT-PCR was performed on a Bio-Rad CFX Manager qRT-PCR instrument (Bio-Rad, America). cDNA was diluted 2-fold (800 ng/uL), and 1 uL was used as template in 20 uL qRT-PCR reaction, with TransStart Top Green qRT-PCR Super Mix (TransGen, China) and gene specific primers qRT-PCR-*TaAFR*-F and qRT-PCR-*TaAFR*-R (S1 Table), and the reaction carried out with an annealing temperature of 58.3°C. A similar reaction was carried out using primers for the wheat reference gene *GAPDH* (GenBank: AF251217) (primers qRT-PCR-*GAPDH*-F and qRT-PCR-*GAPDH*-R, annealing temperature of 58.3°C) (S1 Table). Three technical replicates were conducted for each of three biological replicates per sample. The relative expression of *TaAFR* was evaluated as described by Yu et al. [20]. For samples where a treatment was included, the value of the control treatments were subtracted from those of the treated samples prior to comparing expression with the time zero untreated controls.

#### Subcellular localization

*TaAFR* CDS, minus the stop codon, was inserted upstream of a GFP tag in the pSuper1300 vector (Laboratory preservation), and the recombinant construct was transformed into *Agrobacterium* GV3101. The strain GV3101-pSuper1300-*TaAFR* was injected into *N. benthamiana* leaves at the five-leaf stage, and then observed over a period of 30 to 80 h by fluorescence microscope (Nikon Ti 2, Japan) with an excitation wavelength of 495 nm.

### Identification of TaAFR interacting proteins

#### Yeast-2-hybrid (Y2H)

The *TaAFR* CDS was cloned into the yeast bait vector pGBKT7 which carries the GAL4 DNA-binding domain (BD), and the construct, BD-*TaAFR*, was subsequently transformed into yeast strain Y187. A yeast cDNA library (AD-*cDNA*) previously constructed was used to screen the partner proteins of TaAFR [27]. Y187-BD-*TaAFR* was co-cultured overnight in YPDA media with AH109-AD-*cDNA* at 30°C with gentle agitation (50 r/min). The mated culture was spread onto petri plates with SD-WLHA medium and incubated at 30°C for 3-5 days. Positive clones were sequenced by Beijing Zhongke Xilin Biotechnology Co., Ltd., and the identity of the partner proteins were determined by BLAST alignments.

Once the identities of the positive interaction were determined, the CDS sequences were amplified from cDNA of TcLr15 inoculated with leaf rust strain 05-19-43②. These coding sequences were then inserted into the pGADT7 vector to generate recombinant AD-*Prey* for re-testing the interactions with the bait protein in the BD-*TaAFR* construct. Positive clones were tested by Y2H further detected by β-galactosidase.

To determine which of the TaAFR domain(s) were interacting with the partner proteins, the cDNA sequences of each of the F-box (1-71 aa) and Kelch (72-383 aa) domains of *TaAFR* were inserted into BD vectors. AD-*Prey* that showed positive interactions with BD-*TaAFR* were then screened for interactions with each of these two domains: AD-*Prey* with BD-*TaAFR-F-box* or with BD-*TaAFR-Kelch* were verified by Y2H assay as described above.

#### Bimolecular fluorescence complementation (BiFC)

Y2H positive interactions were validated by BiFC. The CDS of *TaAFR* and the partner proteins were inserted into pSPY CE and pSPY NE vectors (Laboratory preservation) to construct the pSPY CE-*TaAFR* and pSPY NE-*Prey* vectors, respectively. GV3101 with pSPY CE-*TaAFR* and with pSPY NE-*Prey* were combined and co-injected into the *Nicotiana benthamiana* leaves. Fluorescence signal was observed as described in the subcellular localization section.

#### Co-immunoprecipitation (Co-IP)

The positive interactions tested by BiFC were further validated by Co-IP. The CDS of *TaAFR* and the putative partner proteins were inserted into pTF101 (Laboratory preservation) with HA or FLAG tags to construct the recombinant vectors pTF101 HA-*TaAFR* and pTF101 FLAG-*Prey*, respectively. GV3101 containing with pTF101 HA-*TaAFR* and pTF101 FLAG-*Prey* were co-injecting and transiently expressed in *N. benthamiana*. The combination of pTF101 HA-*TaAFR* and pTF101 FLAG-*TaGFP* were used as the negative control. Proteins were extracted from leaves of *N. benthamiana* sampled after co-injecting 60 h and subjected to IP by HA-magnetic beads. The eluted proteins were subjected to immunoblot analysis with anti-FLAG tag polyclonal antibody (Solarbio, China). The detailed Co-IP was performed as described by Zhu and Huq [28].

## Results

### Genome-wide identification of wheat FBKs

#### 74 Wheat FBK proteins were identified and divided into 5 categories based on their functional domains

The seed sequences of F-box (457), F-box-like (306) and Kelch (486) domains were obtained from the Pfam database. “F-box domain” will be used henceforth to describe both F-box and F-box-like domains. A total of 192 transcript sequences containing at least one F-box and/or Kelch domain were identified by searching wheat IWGSC translated transcript database with HMMER3.1b2. Among these, 68 genes encoding 74 transcripts were found to carry both the F-box and Kelch domains predicted by SMART and HMMER. The putative protein sequences (S1 File), their theoretical isoelectric point (*p*I), molecular weight (MW), number of intron, subcellular localization and functional domains of the 74 putative wheat FBKs are presented in S2 Table. Wheat FBK (TaFBK) proteins ranged from 239 to 643 residues in length, with predicted MWs of 27.41-69.41 kDa and theoretical *p*Is of 4.18-9.99, of which the number of acidic/alkaline proteins account for half of the proteins, and 22 of these (29.7%) were greater than 9.0. Most of the TaFBKs were predicted to localize in the nucleus, cytoplasm or plastid, while a handful were predicted to localize in various organelles (peroxisome, golgi). The intron-exon structure has been reported to be closely related to the evolution of the F-box superfamily [29]. The number of introns in 74 *TaFBK* transcripts varied from 0 to 4, among them, 37 *TaFBK* transcripts had 1 intron (50.0%) and 25 transcripts had no intron (33.8%).

Each of the 74 *TaFBK* transcripts contained only one F-box domain at the N-terminus, with up to four Kelch domains at the C-terminus. A few members also carried PAS and PAC domains upstream of the F-box domain. According to their different domain structures, TaFBKs were divided into 5 categories as followed: F-box+1 Kelch, F-box+2 Kelch, F-box+3 Kelch, PAS+F-box+4 Kelch, and PAS+PAC+F-box+4 Kelch (S1 Fig). The F-box+2 Kelch was the largest category, accounting for 40.5%, followed by F-box+1 Kelch (35.1%), while PAS+PAC+F-box+4 Kelch was the least represented with 2 proteins in this group.

#### Wheat FBKs show conservation of F-box and divergence of Kelch domain sequences

MEGA7 was used to align the F-box or Kelch domains of TaFBKs. WebLogo’s were generated, where the height of each stacked letter represents the probability that a given amino acid will be occurred at each position (S2 Fig). In the wheat F-box domain (S2A Fig), L-16 and R-18 are relatively tall, indicated a high probability that those residues would be found at those positions. In the F-box domain alignment, 69 and 65 of the 74 proteins analyzed respectively carried L residues at the 16th position and R residues at the 18th position, which indicates that these 2 amino acids were indeed highly conserved in F-box domain. In addition, L-6 (82.4%), P-7 (81.1%), V-30 (87.8%) and W-34 (81.1%) were fairly conservative, followed by P-20 (73.0%), D-8 (68.9%), R-28(63.5%), R-32 (60.8%), D-9 (56.8%), C-31 (59.5%), V-19 (54.1%), C-15 (52.7%) and A-11 (51.4%).

The Weblogo of the Kelch domain (S2B Fig) showed that G-19 (85.8%), G-20 (86.4%), W-53 (97.9%) and M-59 (54.1%) were highly conserved. In addition, although the height of some residues, such as R-2 (14.11%), H-5 (11.5%), L-10 (15.5%), G-12 (25.0%) and D-45 (20.3%), were shown as relatively high in the Kelch domain, these were poorly conserved according to the statistical assessment. Compared to the other protein sequences, TaFBK65 had 3 additional residues (PVP) at the N-terminus of the F-box motif; these 3 residues were removed in order to prepare the Kelch WebLogo. In general, the amino acid sequences within the Kelch motif were more divergent than that observed within the F-box domain.

#### Phylogenetic distribution of the wheat, Arabidopsis, rice, sorghum and maize FBK subfamilies grouped according to the number of Kelch domains

To understand the evolutionary relationship of the 74 TaFBKs members, a phylogenetic tree was constructed together with 94 Arabidopsis FBKs (AtFBKs), 31 rice FBKs (OsFBKs), 34 sorghum FBKs (SbFBKs) and 32 maize FBKs (ZmFBKs) (Fig 1). The tree resolved into 7 clades, where the AtFBKs were mainly distributed in clade G, all of TaFBKs, OsFBKs, SbFBKs and ZmFBKs, with the exception of three OsFBKs (22, 14 and 9) and three SbFBKs (23, 16 and 8), distributed in clades A to F. All members in clade C belong to the F-box+1 Kelch type, and among them, only 5 members were from Arabidopsis, while the remaining 37 were from Gramineae. In the clades D and F, the FBKs of F-box+2 Kelch type accounted for the largest size, containing only a few members of F-box+1 Kelch and F-box+3 Kelch types. Clade E mainly contains F-box+3 Kelch type FBKs from Gramineae and 3 members of F-box+2 Kelch that come from Arabidopsis. FBKs with 4 Kelch domains (F-box+4 Kelch, PAS+F-box+4 Kelch, PAC+F-box+4 Kelch, and PAS+PAC+F-box+4 Kelch) were from 5 species and absolutely grouped in clade B. AtFBK54 (LSM14+F-box+2 Kelch) grouped with other members of Arabidopsis F-box+2 Kelch in clade G, OsFBK31 and OsFBK28 (F-box+1 Kelch+RING) were divided into clade D. The phylogenetic analysis showed that the number of Kelch domains was a key classification criterion within the FBK subfamily.

**Fig 1.**
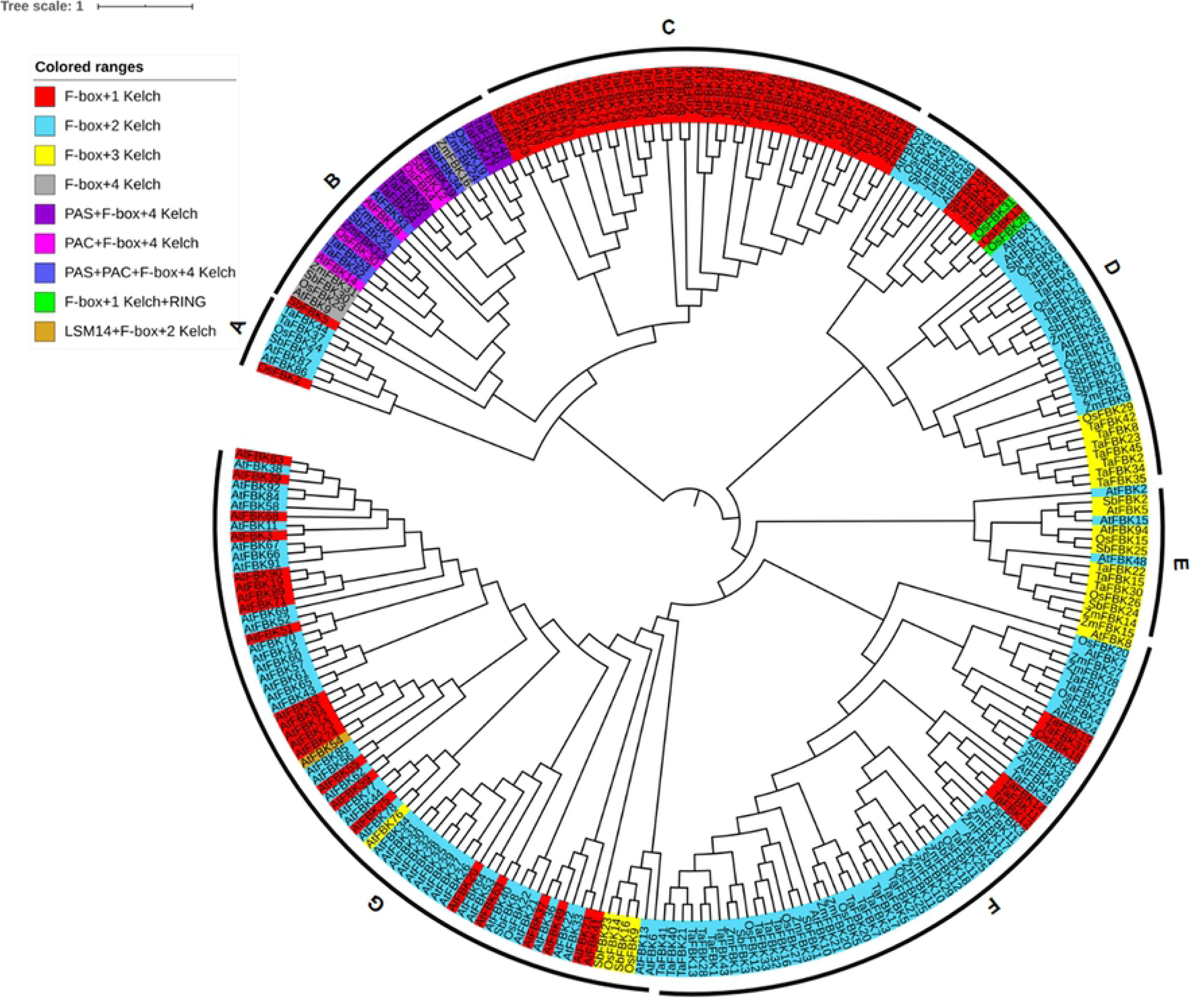
Phylogenetic analysis of FBK proteins in wheat, Arabidopsis and three important monocots. The full-length amino acid sequences were aligned by ClustalX 2.0 and the Maximum Likelihood (ML) tree was constructed using MEGA7. FBK proteins were grouped into 7 distinct clades named A-G.

#### *TaFBK* genes are unevenly distributed on the wheat chromosomes and mainly expanded its size by segmental duplications

The chromosomal position of 68 *TaFBK* genes were retrieved from the EnsemblPlants and a chromosomal distribution map was generated (S3 Fig). The *TaFBK* genes were unevenly distributed on the wheat 21 chromosomes. The chromosomes of 4A (5), 4B (5), 6A (6), 6B (8), and 6D (6) had relatively higher distribution densities, whereas only one *TaFBK* gene was found on each of chromosomes 1B, 3B and 1D, and none were detected on chromosome 2B.

In animals, the number of F-box proteins is relatively low compared with plants, with only 68 and 74 F-box genes in the human and mouse genomes, respectively [30]. Incidentally, the wheat genome encodes the same number of the Kelch subfamily proteins, which represents only a portion of F-box proteins encoded in this species. Gene duplication is thought to be the main driving factor in the expansion of F-box family in plants [17]. To explore the evolutionary mechanism of wheat FBK subfamily, the present study investigated tandem duplication and segmental duplication events in the wheat FBK subfamily of the F-box family by observing similarities among 68 FBK sequences. A total of 57 *TaFBKs* were identified to be segmental or, to a lesser extent, tandem duplications. Among the segmental duplication genes, which were distributed to 20 wheat chromosomes, 8 groups consisted of a pair of genes, 6 groups contained 3 genes, and there were 2 groups each with 6 and 8 genes.

Tandemly duplicated genes affected 8 *TaFBK* genes, and each of these occurred on the 4^th^ chromosomes. These results indicate that both segmental and tandem duplications played a role in the expansion of the *TaFBK* subfamily, and unlike the results of Xu et al. in Arabidopsis and rice species, segmental duplications were more prolific in wheat [6].

#### Tissue-specific and abiotic stress response *in silico* expression of *TaFBKs*

To gleam insights into the putative functions of the identified wheat FBKs, *in silico* expression analysis of these genes was evaluated in different wheat tissues at different developmental stages, and in wheat leaves in response to environmental stresses. The FPKM values of *TaFBKs* from five different tissues and three stress combinations were downloaded from digital PCR data available in WheatExp (S3 Table) and were used to construct a heat map using the zero to one normalized scale method. Tissue-specific expression data (cultivar Chinese Spring) was available for 47 *TaFBKs* (Fig 2). In general, *TaFBK* genes exhibited differential expression in all five wheat tissues, suggesting that these genes may be involved in the developmental regulation of multiple tissues. There were two conditions where tissue-specific expression at specific developmental stages showed significantly less transcript accumulation; these are leaf (z10) and grain (z75). Meanwhile, most *TaFBK* genes were generally more abundantly expressed in the spikes (z32, z39, z65) and grains (z71, z85). *TaFBK3* transcripts specifically accumulated in root tissues, *TaFBK8* and *TaFBK29* expressed dominantly in mature leaf (z71), while *TaFBK60* and *TaFBK61* showed highest expression in root tissues followed by grain samples.

**Fig 2.**
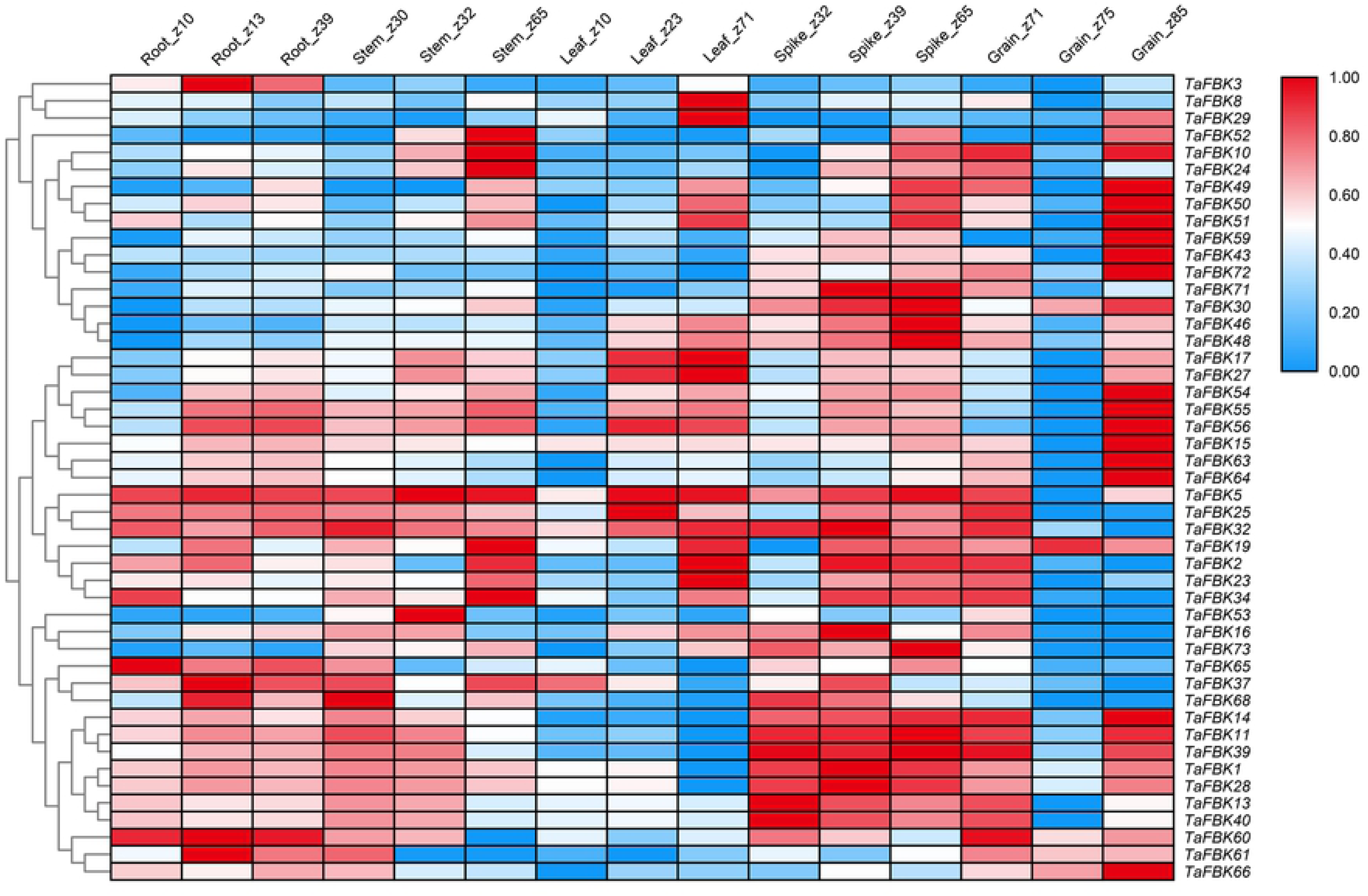
Heat map showing digital expression profiles of *FBK* genes in various tissues and at different developmental stages of wheat based on FPKM values. The color key represents FPKM values. Identity of tissue samples and developmental stages (Zadoks scale) are provided at the top of each lane.

A second data set from the WheatExp database was analyzed for the effect of drought (DS), heat (HS) and heat+drought (HD) stresses on the expression of the same 47 *TaFBK* genes in seedlings of the wheat cultivar TAM 107. A heat map was generated for this dataset showing differential expression at 1 h and 6 h (Fig 3). A general overview of expression of *TaFBK* genes affected by DS was as follows: 34.0% of the *TaFBK* genes were up-regulated; 47.0% genes showed strongly or slightly down-regulated expression; and 19.0% (9 transcript) maintained stable expression between treatment and control. Following heat treatment (HS) at 40°C, the following changes were observed: transcripts *TaFBK60*, *TaFBK61* and *TaFBK*46 increased sharply at 1 h, and then decreased at 6 h; *TaFBK10*, *TaFBK23* and *TaFBK*50 expression gradually increased from 0 h (control) to 6 h; eight *TaFBKs* were down-regulated at both time points assessed; transcripts of seven genes (14.9%) decreased to the roughly half of the control levels at 1 h; expression of the remaining 38.3% (18) genes sharply declined at 1 h after the stress treatment, and transcript accumulation of 7 of these 18 genes returned to levels similar to that of the control by 6 h, while the remaining 9 genes increased slightly at 6 h compared with the earlier time point. Following the combined treatment HD: 25.5% of the genes increased their expression from 1 h to 6 h compared with control; transcripts from seven genes gradually decreased from 1 to 6 h; 72.3% transcripts sharply decreased at 1 h treatment, then slightly or sharply increased at 6 h. In brief, HS caused more obvious and intense change on expression of *TaFBKs* when comparing to DS treatment.

**Fig 3.**
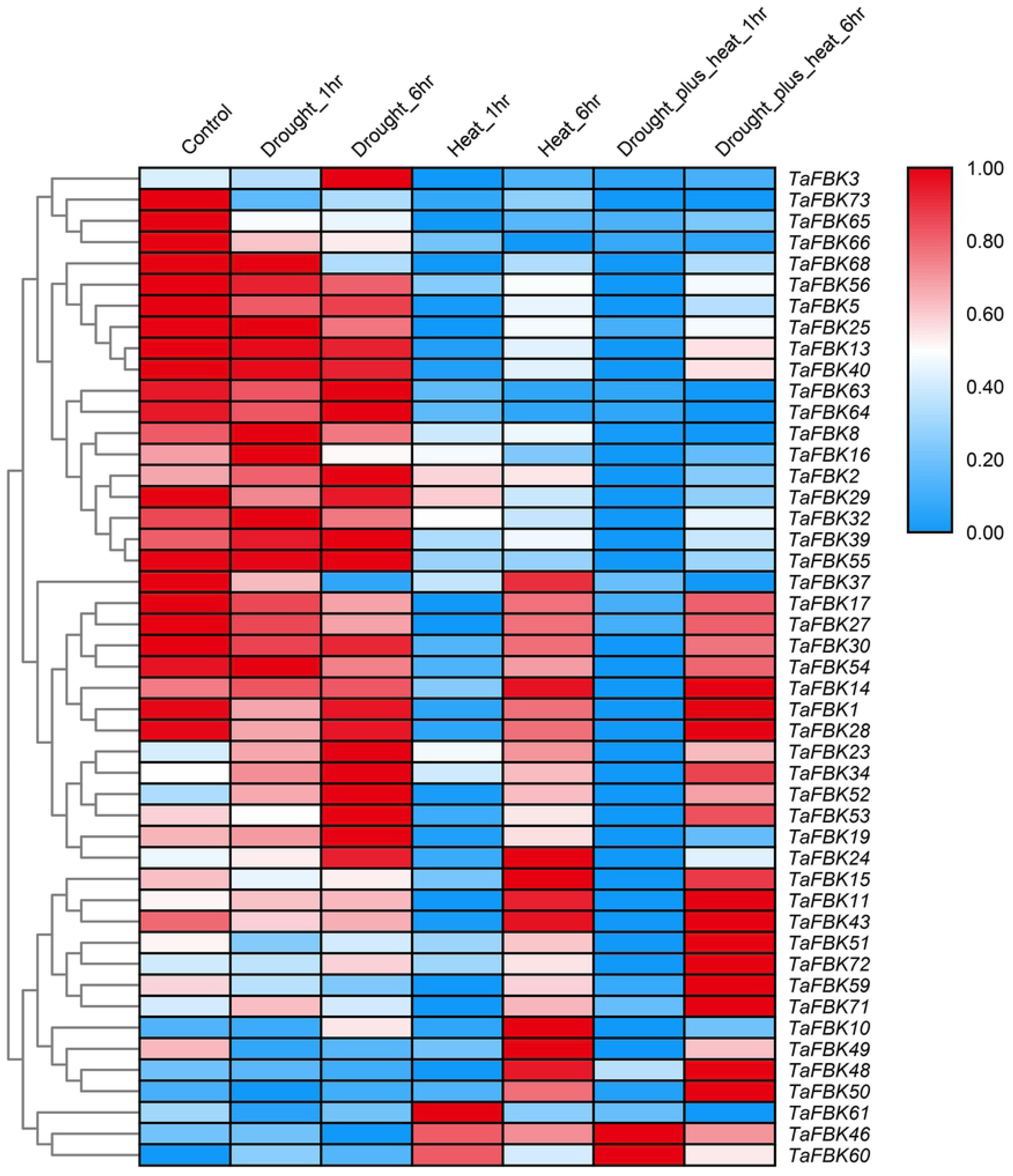
Heat map showing digital expression profiles of FBK genes in wheat response to DS, HS and HD based on FPKM values. Color key represents FPKM values. The method and time of treatments are provided at the top of each lane. DS, drought stress; HS, heat stress; HD, heat+drought stresses.

### Molecular identification and expression patterns of *TaAFR*

#### *TaAFR* gene encodes a wheat FBK protein

The heat map presented in Fig 3 showed that the expression of *TaFBK19* was strongly up-regulated under DS treatment and sharply down-regulated under HS treatment at 1 h, while the combined heat and drought (HD) treatment resulted in *TaFBK19* in low level from 1 to 6 h. This gene was selected for further investigation of its expression pattern by qRT-PCR. At first, we cloned the full-length (1327 bp) cDNA sequence from TcLr15 wheat seedlings inoculated with the leaf rust strain 05-19-43②. The cDNA encodes a polypeptide with 383 amino acids. The MW of the predicted polypeptide was 40.69 kDa, and the predicted *p*I was 5.11. BLASTx analysis showed the sequence shared very high similarity (94%) with an F-box protein, AFR-like, from *Aegilops tauschii* (GenBank: XP 020194469.1). TaBFK19 protein carries a single highly conserved F-box domain (32-71 aa sites) at the N-terminus and a fairly divergent Kelch domain (136-174 aa sites) at the middle region (Fig 4A). Phylogenetic analysis indicated that the TaFBK19 protein shared 94.10% and 87.47% similarity with AFR from *A. tauschii* and *Hordeum vulgare*, respectively, followed by AFR from *Brachypodium distachyon*, *O. sativa*, *Setaria italica*, *Panicum hallii*, *S. bicolor* and *Zea mays*. Meanwhile, AFRs from woody plants (*Prunus avium*, *Musa acuminata*, *Elaeis guineensis*, *Phoenix dactylifera*) and dicots (*Nelumbo nucifera*, *Dendrobium catenatum* and *A. thaliana*) were grouped in different clades (Fig 4C), which indicates that these FBKs were conserved in monocots. Based on the similarities between *TaFBK19* and *AFR* genes from the cereal and monocot, *TaFBK19* will henceforth be described as *TaAFR*. Sequence analyses of TaAFR did not reveal any predicted signal peptide or transmembrane domains, and the protein is predicted to localize to the cytosol. The predicted 3D structure showed three distinct α-helices at the N-terminus and β-sheets at the C-terminal end. The β-sheets are predicted to form 6 triangles, which further cluster to a regular hexagonal arrangement. These secondary and ultra-secondary structures indicated that the protein folds into chair-like configuration (Fig 4B).

**Fig 4.**
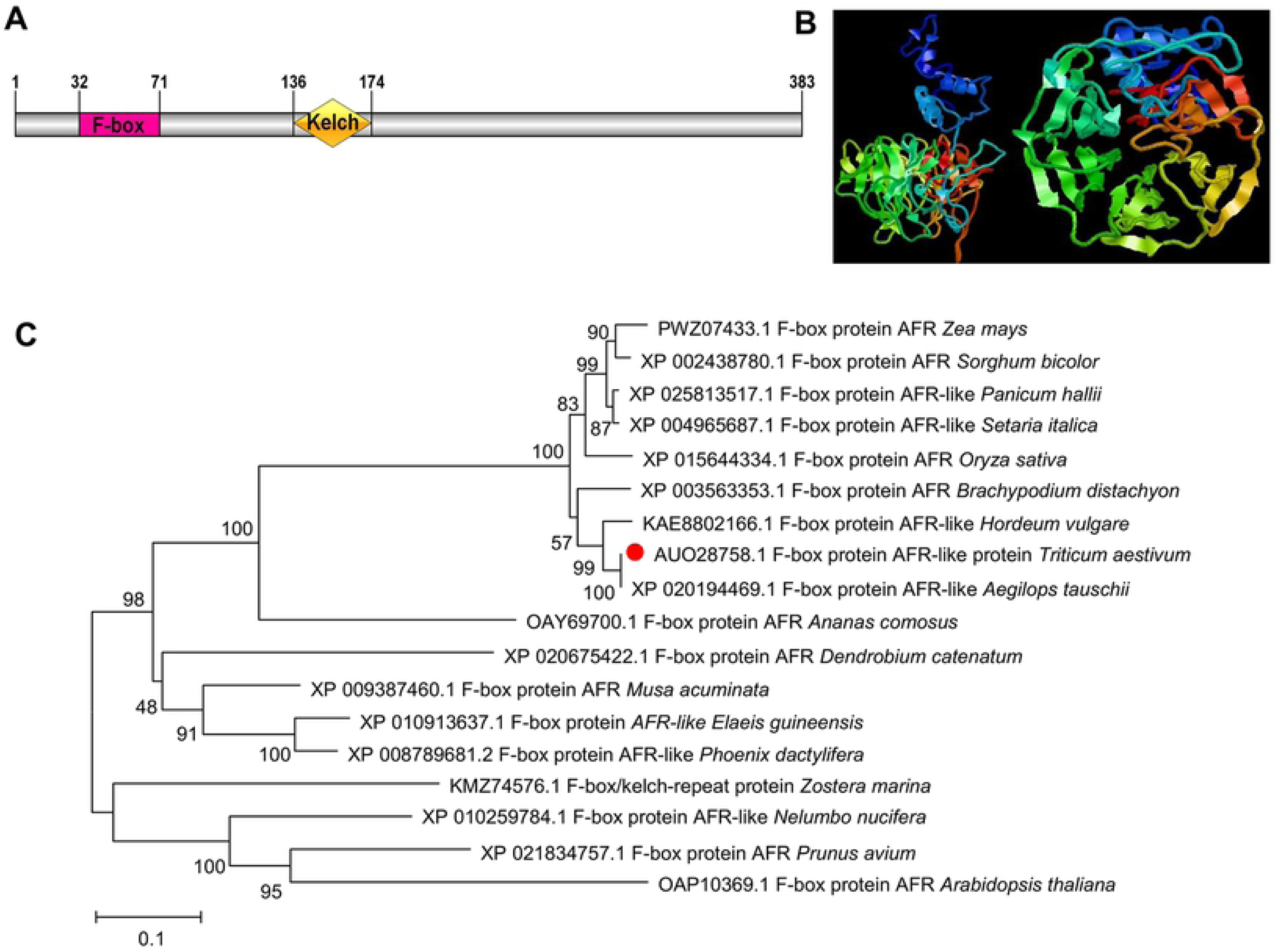
Sequence characteristics of wheat TaAFR. (A) Functional domain of TaAFR. A schematic diagram showing the positions of F-box domain and Kelch domain in TaAFR; (B) 3D structure prediction of the TaAFR protein. The blue helix represented the α-helix structure, and the arrow represented the β-sheet structure. Blue and red differentiate the N- and the C-terminus, respectively; (C) Phylogenetic analysis of TaAFR with F-box proteins from different plants species. The phylogenetic tree was generated using the neighbour-joining method in MEGA 7. Branches were labeled with the GenBank accession number followed by species name.

#### *TaAFR* is primarily expressed in wheat leaves

Six tissues were sampled from wheat TcLr15 seedlings (root, leaf and stem) and adult plants (pistil, stamen, flag leaf) to analyze the tissue-specific expression of *TaAFR*. The young leaf was used as a control (the expression value was set 1.0) to measure it’s relative expression to other tissues. *TaAFR* was mainly expressed in young leaf, with lower expression in the flag leaf and extremely low expression was detected in young root, pistil, and stamen (Fig 5A).

**Fig 5.**
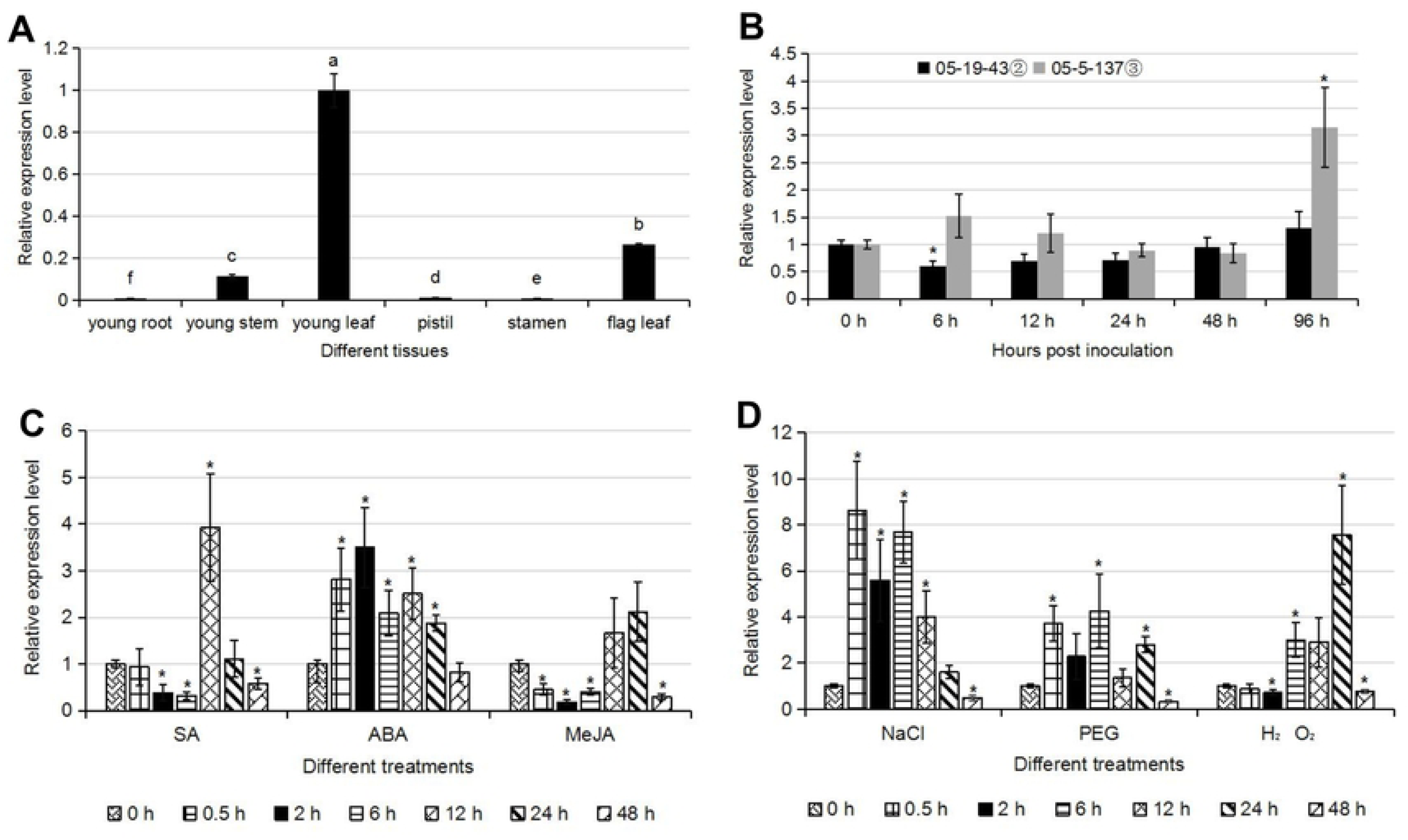
The expression patterns of *TaAFR* in different wheat tissues and stress/hormone treatments. (A) Expression profile of *TaAFR* in different tissues of TcLr15; (B) Expression patterns of *TaAFR* gene in incompatible and compatible combinations of TcLr15/*Puccinia triticina* strains 05-19-43② and 05-5-137③; (C) Effects of SA, ABA and MeJA on expression of *TaAFR* in TcLr15 leaves; (D) Effects of NaCl, PEG and H_2_O_2_ on expression of *TaAFR* in TcLr15 leaves. The control leaf of TcLr15 were sampled at the corresponding time points, and these samples was used as the subject to be subtracted. Different letters indicate significant differences (p < 0.05) for tissue-specific comparisons; An asterisk, *, marks the significant difference between treatment and the 0 h untreated control (p < 0.05).

#### Differential expression of *TaAFR* in compatible and incompatible wheat/leaf rust pathogen combinations

After inoculation with different virulent leaf rust strains, the temporal expression profile of *TaAFR* in TcLr15 is shown in Fig 5B. Generally, the *TaAFR* transcript was higher in the compatible interaction (TcLr15 inoculated with 05-5-137③) than in the incompatible one (TcLr15 inoculated with 05-19-43②) except 48 hpi. For the incompatible interaction, the *TaAFR* transcripts gradually increased from 6 to 96 hpi, but apart from the initial increase from 0 to 6 h, no significant difference was observed across the time course. In the compatible interaction, no significant change was observed between 0 to 48 hpi, but a rapid increase was observed at 96 hpi, where the expression of *TaAFR* transcripts was 3.1-fold higher than in the 0 h untreated control samples.

#### Exogenous SA and ABA applications significantly up-regulated the expression of *TaAFR*

The expression pattern of *TaAFR* in TcLr15 following exogenous treatment with plant hormones is presented in Fig 5C. In response to SA treatments, *TaAFR* expression was down-regulated 2-fold at 2 and 6 h compared with the untreated control, and thereafter increased rapidly 4-fold at 12 h compared with the control before returning to the basal expression levels. In response to ABA, *TaAFR* expression increased rapidly by 3.5 folds, observed at 2 h, and continued to be up-regulated throughout the time course, decreasing gradually until 48 h where basal level expression was observed. MeJA application resulted in significant down-regulation of *TaAFR* at most time points, except at 12 and 24 h, where no significant difference was observed compared with the control.

#### Expression of *TaAFR* is affected by salt, drought and oxidative stresses

Three abiotic stress treatments, salt (NaCl), drought (PEG) and oxidative stress (H_2_O_2_), were evaluated for their effects on *TaAFR* expression in TcLr15 seedlings. The expression of *TaAFR* was significantly affected in TcLr15 after treatment with NaCl (Fig 5D). The transcripts were strongly up-regulated from 0.5 h, and maintained a high level of expression until 12 h. Two expression peaks occurred at 0.5 h (8.2-fold) and 6 h (7.9-fold), respectively. Thereafter, the *TaAFR* transcripts started to down-regulate gradually, until 48 h, where transcripts dropped to half of the level detected in the 0 h untreated controls. In response to PEG treatments, the *TaAFR* transcript was increased in abundance at 0.5 h (3.9-fold), 2 h (2.1-fold), 6 h (4.1-fold) and 24 h (3-fold), but was down-regulated at 48 h. After treatment with H_2_O_2_, the expression of *TaAFR* did not differ from that of the control until 2 h after treatment when it was down-regulated, but from 6 h to 24 h, *TaAFR* showed an upward trend, reaching a peak at 24 h where it was 7.8-fold higher than that of the 0 h control. Finally, expression dropped below the 0 h control levels at 48 h (Fig 5D).

#### TaAFR is localized to the nucleus and cytoplasm

*Nicotiana benthamiana* was injected with GV3101 containing either the empty vector 35S:*GFP* or the recombinant vector 35S:*TaAFR*-*GFP*, and transient expression of the recombinant proteins was observed. The fluorescence signal of 35S:*GFP* was visualised in both the nucleus and cytoplasm after 36 h transfection (Fig 6), whereas the fluorescence signal of 35S:*TaAFR*-*GFP* was detected after 48 h, predominantly observed in the nucleus and cytoplasm. Moreover, the nuclear dye DAPI was used to stain the tobacco leaves after transfection, light blue was clearly observed in the nucleus.

**Fig 6.**
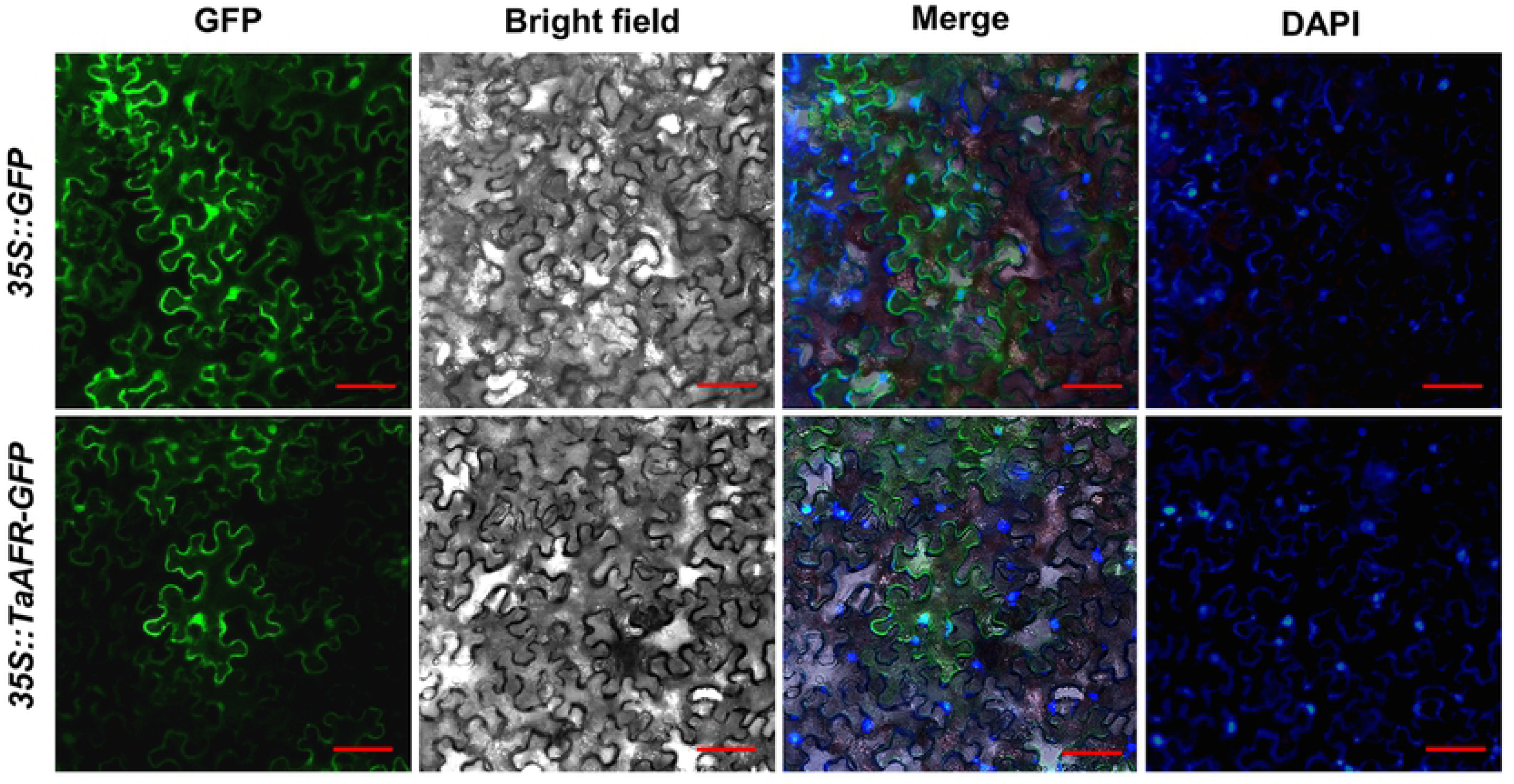
Fluoresence observation for subcellular localization of TaAFR. The free GFP protein and TaAFR-GFP fusion protein were transiently expressed in the *N. benthamiana* by *Agtobacterium*-mediated transformation. GFP, GFP fluorescent signal channel; Bright field, ordinary light channel; DAPI, nuclei were stained by DAPI; Merge, merge of GFP, Bright field and DAPI. Bar=20 μm.

### Screening and identifying the partner proteins interacting with TaAFR

#### Thirteen types of proteins putatively interacted with TaAFR

To identify candidate upstream and/or downstream proteins interacting with TaAFR in wheat, we screened a yeast library carrying the cDNA of TcLr15 inoculated with incompatible leaf rust strain against the bait construct, BD-*TaAFR.* Clones from positive interactions were sequenced and thirteen candidate proteins were identified from 47 clones. Candidate proteins are listed in S4 Table, and categorized into the following 5 groups: photosynthesis, stress resistance, transportation, basal metabolism and unknown protein. Among these, 5 stress resistance related proteins were obtained: such as Peroxidase 51-like (POD), obtusifoliol 14-alpha-demethylase (CYP51), Glucan endo-1,3-beta-glucosidase 14 (GV), Laccase-7 (Lac7) and leucine-rich repeat protein 1 (LRR-8 Superfamily) (LRR) [31–35]. Meanwhile, transport related proteins, ADP-ribosylation factor 2-like isoform X1 (ARL2) and SEC1 family transport protein SLY1 (SLY1) and basal metabolism related protein Skp1/ASK1-like protein (Skp1) were also detected [36, 37, 3].

#### TaSkp1, TaARL2 and TaPAL interacted with TaAFR

Based on the results of Y2H library screening, we obtained the complete coding region of Rubisco, Skp1, ARL2, GV, RP, SLY1, NADH, POD, LRR, Lac7 and CYP51, from TcLr15 for further validation of protein interactions. According to Zhang et al., Kelch repeat F-box proteins are regulated by phenylpropanoid biosynthesis by controlling the turnover of phenylalanine ammonia-lyase (PAL) [11]; therefore, in addition to the positive interactions identified in the Y2H assay, we isolated a *PAL* gene from TcLr15. Basic characteristics of these proteins are presented in S5 Table. The interactions were first re-verified by Y2H. Colonies with blue pigments are indicative of positive interactions, and along with the positive control, six such interactions were observed: BD-*TaAFR* and AD-*TaSkp1*, BD-*TaAFR* and AD-*TaSLY1*, BD-*TaAFR* and AD-*TaARL2*, BD-*TaAFR* and AD-*TaCYP51*, BD-*TaAFR* and AD-*TaPAL*, BD-*TaAFR* and AD-*TaNADH*. The remaining combination, along with the negative control, did not grow on the SD-WHLA plates. These results suggest TaAFR might physically interact with TaSkp1, TaSLY1, TaARL2, TaCYP51, TaPAL and TaNADH (Fig 7).

**Fig 7.**
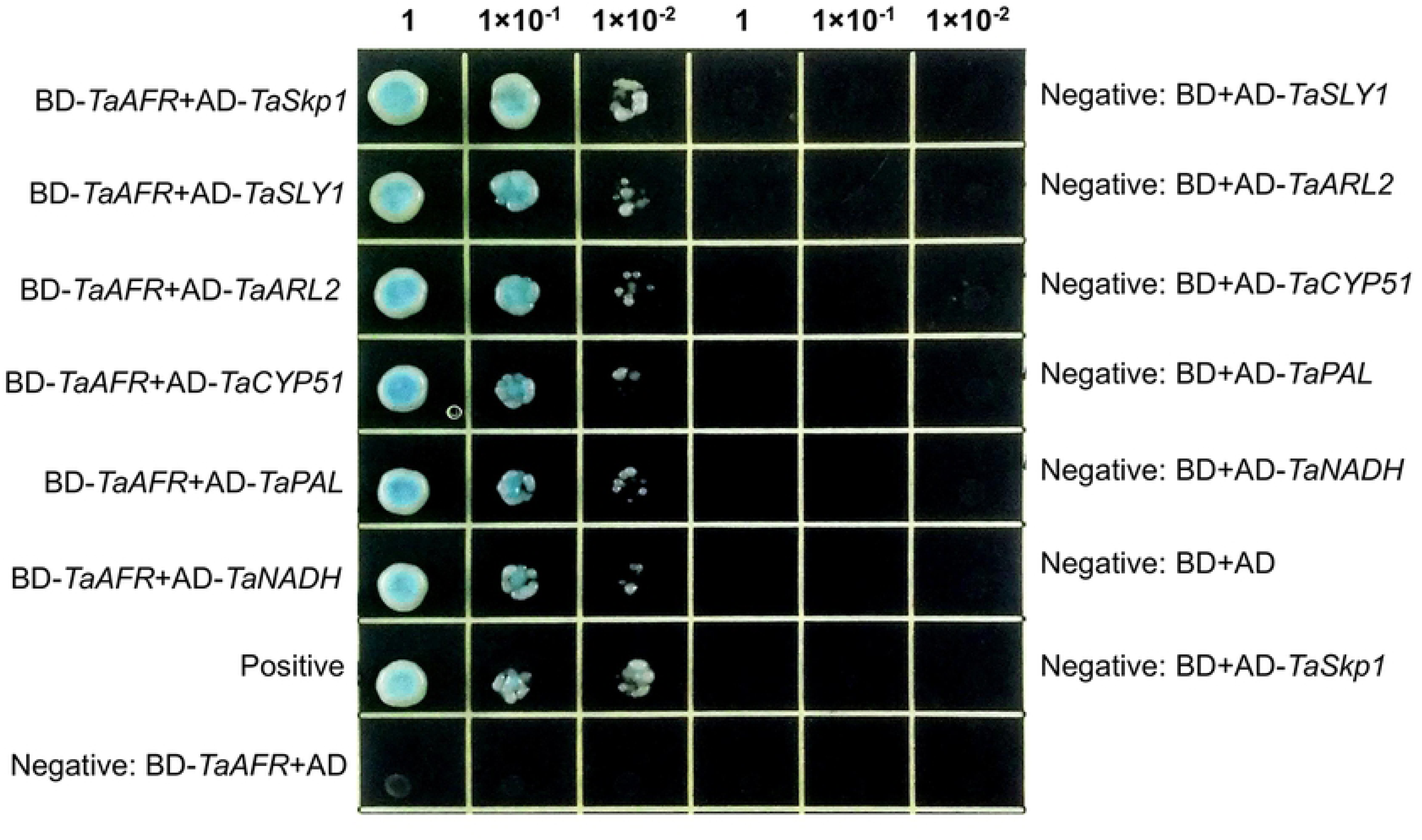
Protein interactions tested by Y2H. Yeast was cultivated on SD-WLHA+X-α-Gal plates for 3-5 days.

To further validate the above results, these 6 interactions were tested by BiFC. In this approach, the coding region of the *TaAFR* was inserted downstream of the c-Myc tag of pSPY CE vector (pSPY CE-*TaAFR*); meanwhile the ORFs of *TaSkp1*, *TaSLY1*, *TaARL2*, *TaCYP51*, *TaPAL* and *TaNADH* were inserted downstream of the 35S promoter in the pSPY NE vector (pSPY NE-*Prey*). The pSPY CE-*TaAFR* vector was used co-transfection of tobacco leaves with each of the pSPY NE-*Prey* constructs. If the gene products of the two constructs interact, a fluorescence signal will be emitted. Among the six combinations, three were found to emit fluorescent signals (Fig 8). pSPY CE-*TaAFR* and pSPY NE-*TaSkp1* emitted fluorescent signal in the nucleus and cytoplasm 40 h after injection. The pSPY CE-*TaAFR* and pSPY NE-*TaARL2* emitted a strong signal in the nucleus and cytoplasm 48 h after co-transfection. The pSPY CE-*TaAFR* and pSPY NE-*PAL* interaction was observed in the cytoplasm by complementary chimeric fluorescence signals 40 h after co-transfection. Thus, the BiFC assay further validated TaAFR interactions with TaSkp1, TaARL2, and TaPAL.

**Fig 8.**
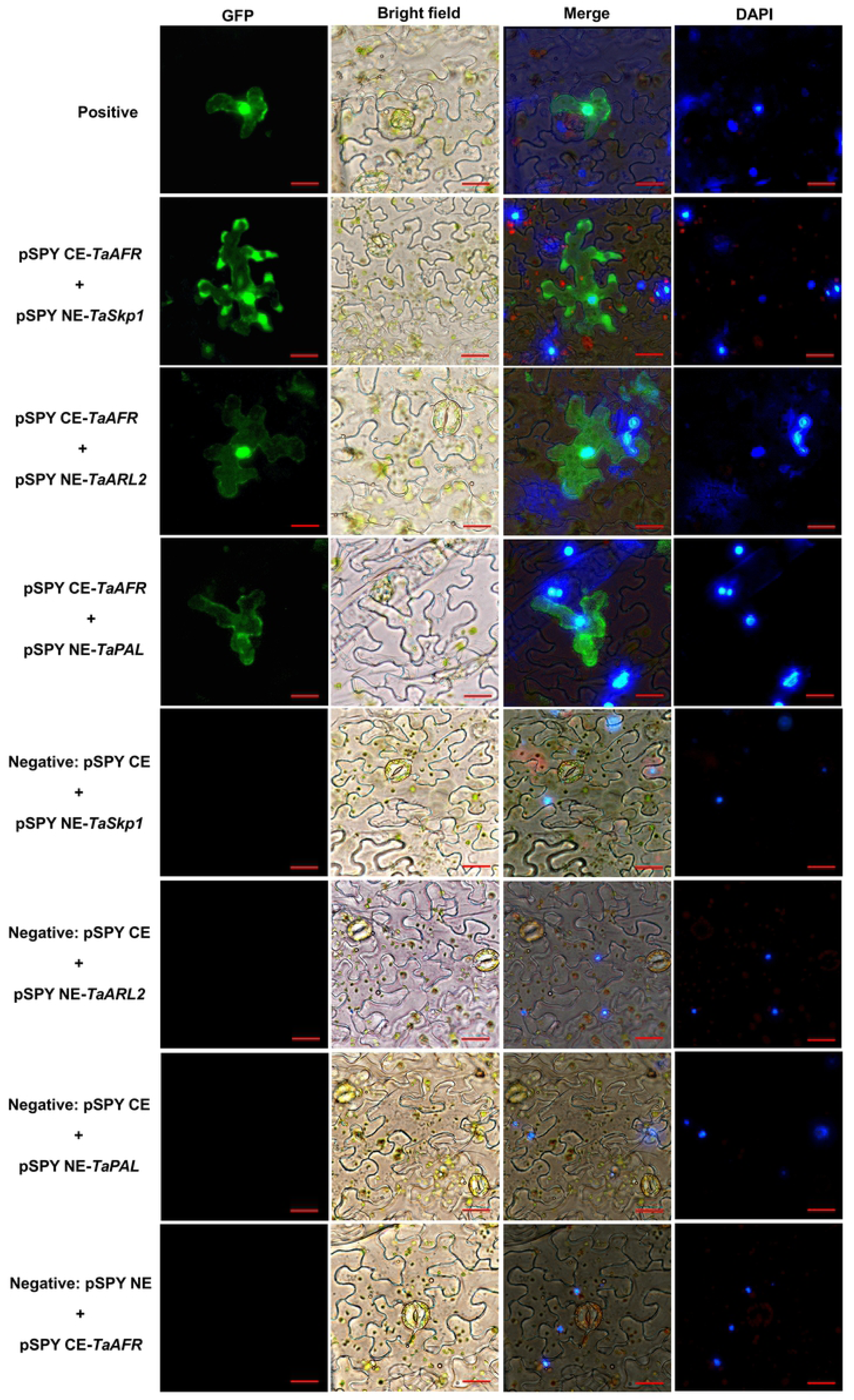
Verification of protein interactions by BiFC. The fluorescence microscope (Nikon Ti 2, Japan) with an excitation wavelength of 495 nm was used to observe fluorescence signal. Three independent experiments were conducted for each combination. Bar= 20 μm.

Co-IP assays were performed upon transient expression in *N. benthaminana* to further validate the results tested by Y2H and BiFC *in vivo*. The combinations of TaAFR with three putative partner proteins TaSkp1, TaARL2, TaPAL and negtive control GFP were successfully detected in the whole cell lysates (WCL). After IP by HA-magnetic beads, the eluted proteins were subjected to immunoblot analysis with anti-FLAG antibody, we found that TaSkp1, TaARL2 and TaPAL were immunoprecipitated with TaAFR since a single band appeared at their corresponding MW sites, but no band except HC (IgG heavy chain) was detected in the combination of TaAFR and GFP (Fig 9). Taken together, these observations support that TaAFR interact with TaSkp1, TaARL2 and TaPAL *in vivo*.

**Fig 9.**
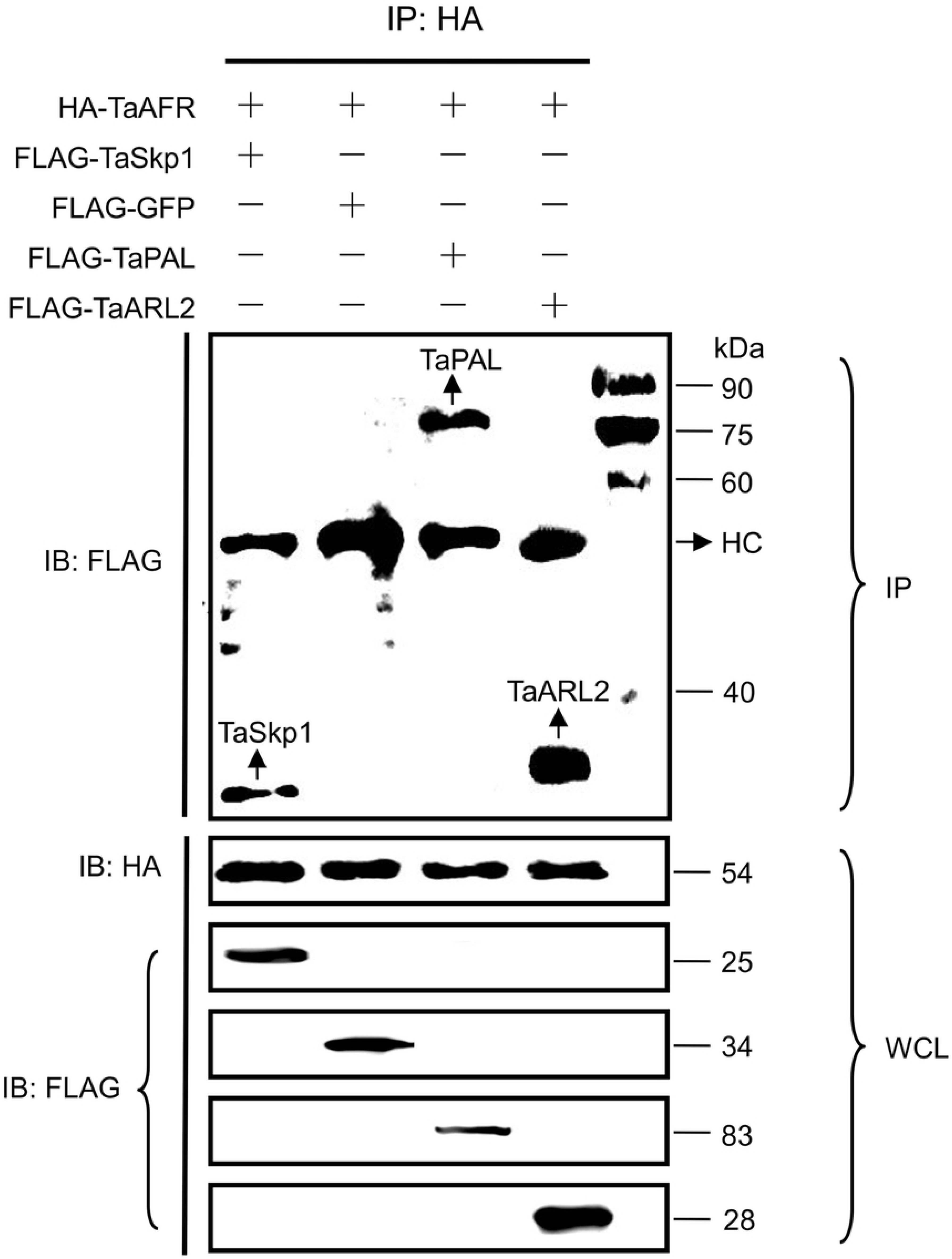
Verification of protein interactions by Co-IP. Proteins were extracted from leaves of *N. benthamiana* 60 h co-injection, and immunoblotting (IB) was used to detect the expression of TaAFR, TaSkp1, TaARL2, TaPAL and GFP in the whole cell lysates (WCL) with HA or FLAG antibody, these proteins were immunoprecipitated by HA-magnetic beads, then the eluted proteins were subjected to IB analysis with anti-FLAG antibody. HC: IgG heavy chain. Marker: 25-90 kDa.

#### The F-box domain of TaAFR interacted with TaSkp1, and the Kelch domain with TaARL2 and TaPAL

To determine which domain of TaAFR is responsible for recognizing the TaSkp1, TaARL2 and TaPAL, we further obtained the cDNA sequences of F-box domain (1-71 aa) and Kelch (72-383 aa) domain of TaAFR, then constructed recombinant BD vector for each. Six combinations of AD-*TaSkp1* and BD-*TaAFR-F-box*, AD-*TaSkp1* and BD-*TaAFR-Kelch*, AD-*TaARL2* and BD-*TaAFR-F-box*, AD-*TaARL2* and BD-*TaAFR-Kelch*, AD-*TaPAL* and BD-*TaAFR-F-box*, AD-*TaPAL* and BD-*TaAFR-Kelch* were verified using the Y2H assay. Among them, three combinations of AD-*TaSkp1* and BD-*TaAFR-F-box*, AD-*TaARL2* and BD-*TaAFR-Kelch*, AD-*TaPAL* and BD-*TaAFR-Kelch* grew well on the SD-WLHA+X-α-Gal plates (Fig 10). These results indicated that TaSkp1 interacted with the F-box domain, while TaARL2 and TaPAL were recognized by the Kelch domain of TaAFR.

**Fig 10.**
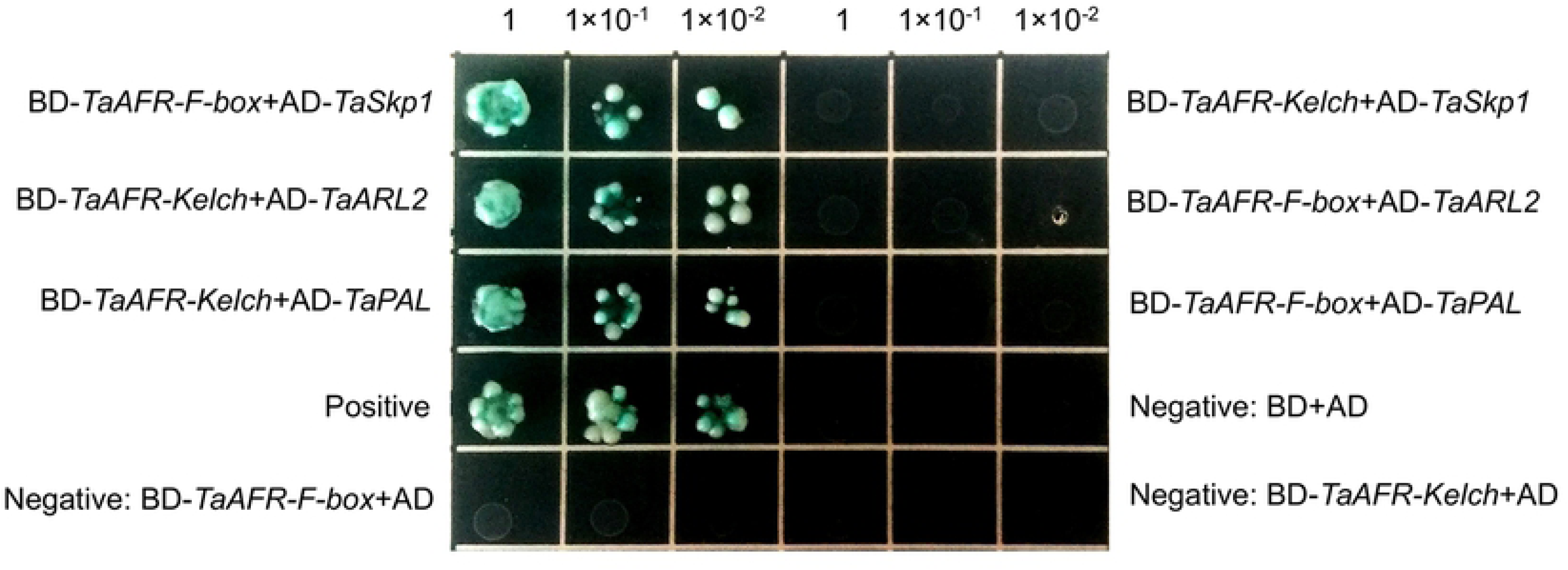
Domain interactions tested by Y2H. Yeast was cultivated on SD-WLHA+X-α-Gal plates for 3-5 days.

## Discussion

In plants, the Kelch type F-box protein is one of the most common subfamilies in the F-box family [38]. Many wheat databases are being continuously updated, with improved annotations over recent years, making it possible for genome-wide identification and comprehensive analysis of gene families. In 2020, Hong et al. reported 41 wheat F-box/Kelch genes, and our previous result from searching against Phytozome 12 (v2.2) database identified 59 wheat F-box/Kelch genes [39, 40]. In the present study, we screened latest IWGSC database in EnsemblPlants with more seed sequences and identified 68 *TaFBK*s encoding 74 putative proteins. The TaFBK subfamily was divided into 7 categories based on differences in number of the Kelch domains. Most of the AtFBKs resolved to clade G and showed relatively distant evolutionary relationship compared with the four Gramineae species, while TaFBKs grouped into the same clade or closer to OsFBKs, SbFBKs and ZmFBKs according to their construction of their functional domains. Compared with the Arabidopsis FBK subfamily, three types of FBKs, namely F-box+4 Kelch, PAC+F-box+4 Kelch and LSM14+F-box+2 Kelch, were not detected in wheat. Each of these types are poorly represented in Arabidopsis, with only one member for each. It may be that they are absent in wheat due to selective evolution of the species, or it may simply be that they cannot yet be detected at the current sequencing depth or annotation of the wheat protein database. F-box+1 Kelch+RING and LSM14+F-box+2 Kelch were found to be unique FBK types in rice and Arabidopsis, respectively.

Studies have shown that FBKs can be localized to the nucleus, cytoplasm and/or organelles. For example, CarF-Box1 (chickpeas) and TML (legume) were localized to the nucleus, while TaKFB1-TaKFB5 (colored wheat) were each co-localized to both the nucleus and cytoplasm [41, 42, 39]. The wheat FBKs identified herein, were predicted to localize in the nucleus, cytoplasm, plastid and/or other organelles. These predictions may provide some insights into potential gene function, but may not always be accurate. For example, the wheat FBK, TaAFR (TaFBK19), was predicted to localize in the cytosol, but was shown experimentally herein to localize in both the nucleus and the cytoplasm.

To gleam some insights into the potential functionality of the wheat FBKs, their expression was observed in response to different stresses. An initial *in silico* analysis was carried out by comparing expression of 47 *TaFBKs* in their response to DS, HS, and HD. These 47 genes showed varied expression patterns in response to these differential treatments, most of them showed strong down-regulation in wheat treating with HS. *TaAFR* was further selected to observe its’ expression patterns related with abiotic/biotic stresses and hormones. NaCl and H_2_O_2_ caused most strong up-regulation of *TaAFR*, moreover, *TaAFR* represented different expression patterns in TcLr15 inoculating with virulent/avirulent leaf rust pathogen. Many previous studies have reported similar observations for the expression of different plant *FBKs* in response to different abiotic and biotic stress. For example, the nuclear localized FBK gene *CarF-box1* from chickpea was shown to play an important role in abiotic stress, where expression levels of this gene were significantly up-regulated after drought and salt treatments, but was down-regulated under heat and cold stresses [41]. The grape FBK gene *BIG24.1* was up-regulated by Botrytis infection in grapes, and the up-regulation expression of this gene also affected the plants response to other biotic and abiotic stresses [43]. In another example, the F-box protein containing two Kelch repeats in sugar beet, homologous to Arabidopsis FBK AT1G74510, was found to interact with the beet necrotic yellow vein virus pathogenicity factor P25, and it was speculated that P25 could affect the formation of SCF complex [44].

Biotic and abiotic stress responses are often regulated by plant signaling hormones and exposure to such stresses can activate these pathways [45]. It is therefore interesting that the expression of *TaAFR* was also affected by three different plant hormones. SA, ABA and MeJA treatments had a medium effect on the expression of *TaAFR*, suggesting that this gene may regulate and be regulated by different plant hormones. A regulatory behavior in plant hormone responses would be consistent with various other FBKs in the hormone signaling pathways [22, 24].

FBKs interact both with other members of the UPS and with downstream targets for proteasome degradation; identification of some of these interacting proteins can further provide insight into the function of this protein. A multifaceted approach was employed to identify and validate candidate interactions. First, using a leaf rust pathogen treated TcLr15 wheat leaf cDNA library, a Y2H library screen was utilized as a broad scale approach to fish for candidate interacting proteins. A total of 13 candidates were identified, and 11 of these were cloned and re-screened by Y2H for interactions with TaAFR. Additionally, a *PAL* gene, which was not identified in the pool, but has been shown to be involved in regulation process of FBKs, was added to the list. Among these, a total of 6 interactions, including the TaAFR-TaPAL interaction, were confirmed positives. However, since Y2H assays can pick up false positives, these 6 genes were then validated using the BiFC and Co-IP methods, and finally three partner proteins interacting with TaAFR were confirmed: TaSkp1, TaARL2, and TaPAL.

To further characterize their detailed interacting domain, another Y2H assay was carried out between the F-box and Kelch domains of TaAFR with each of these proteins. TaSkp1 was shown to interact with the F-box domain but not with the Kelch domain. This was not unexpected since Skp1 is a known component of the SCF complex and F-box proteins interact with Skp1 via the F-box domain. This result provides preliminary evidence that TaAFR forms part of the SCF complex. Meanwhile, the other two proteins, TaARL2 and TaPAL, were shown to interact with the Kelch domain, and not the F-box domain, suggesting that these two proteins are targeted by TaAFR for ubiquitination and designated for proteolytic degradation.

ADP ribosylation factor (ARF) family of small GTP binding proteins regulates a wide range of cellular processes in eukaryotes [46]. According to Guan et al., three ARF genes (*PvArf1*, *PvArf-B1C* and *PvArf-related*) were identified and localized in the nucleus and cytoplasm, which regulated proline biosynthesis by physically interacting with PvP5CS1 to improve salt tolerance in Switchgrass (*Panicum virgatum* L.) [47]. The latest research showed that wheat contains 74 *TaARF* genes. The expression of *TaARFA1* genes was regulated by biotic stress (powdery mildew and stripe rust pathogens) and abiotic stress (cold, heat, drought and NaCl), and may be related to the ABA signaling pathway [48]. ARL2 (ADP-ribosylation factor 2-like) is most closely related to the ARL2 subfamily of ARF-like (ARL) proteins. ARL2 localizes in the cytosol, centrosomes, nucleus, and mitochondria [49]. Arabidopsis *TTN5* encodes an ARL protein, and functioned throughout the Arabidopsis life cycle, with an important role in the regulation of intracellular vesicle transport [36]. Most of the previous research of ARL2 had been focused mainly on humans and yeasts, with little information in plants. Thus, future studies on the interaction of TaAFR and TaALR2 may provide valuable insights in the function of both proteins in plants.

PAL activity is modulated by abiotic/biotic stresses in plants, including infections with fungal pathogens, UV/blue light irradiation, and wounding [50]. Zhang et al. found that differential expression of an Arabidopsis *FBK* genes affected the stability of PAL, and PAL isozymes were shown to physically interact with FBKs both *in vitro* and *in vivo* [11]. The interaction of PAL with FBKs thereby controls phenylpropanoid biosynthesis by mediating the ubiquitination and subsequent degradation of PAL. In another study, the authors showed that the Arabidopsis FBK protein, KFB39, a homolog of AtKFB50, also interacted with PAL isozymes and regulated PAL stability and activity, thereby participating in the plant’s tolerance to UV irradiation [12]. In the work presented herein, TaAFR interacted with PAL, presumably through the Kelch domain which was also shown to interact with PAL. These results, together with the observations of Arabidopsis FBK activity in Zhang et al., point to the possibility that TaAFR regulates PAL stability and activity in the wheat response to abiotic/biotic stresses [11, 12]. Meanwhile, the work presented in this manuscript provides a glimpse into their potential function, and opens the door for future studies to further characterize these genes.

## Conclusion

A total of 68 *TaFBK* genes encoding for 74 proteins were identified in wheat in a genome-wide survey. The FBK proteins from wheat, Arabidopsis and three important monocots were grouped into 7 clades according to the number of Kelch domain. 68 *TaFBK* genes were unevenly distributed on 21 wheat chromosomes, *TaFBKs* differentially expressed at multiple developmental stages and tissues, and in response to drought and/or heat stresses by *in silico* analysis. A Kelch type F-box gene *TaAFR* was isolated and identified to localize in the nucleus and cytoplasm, which primarily expressed in wheat leaves, and also revealed varied expression patterns in response to treatments with leaf rust pathogens, exogenous hormones, and abiotic stresses. Skp1 interacted with the F-box domain of TaAFR, while ARL2 and PAL were recognized by Kelch domain. This work provides a foundation from which to build more detailed research inquiries into the function of the numerous wheat FBKs and also to further characterize the *TaAFR* gene.

## Author contributions

**Conceptualization:** Chunru Wei, Xiumei Yu.

**Data curation:** Chunru Wei, Runqiao Fan.

**Investigation:** Chunru Wei, Yuyu Meng.

**Software:** Chunru Wei.

**Writing - original draft preparation:** Chunru Wei.

**Methodology:** Yiming Yang, Xiaodong Wang.

**Writing - review & editing:** Weiquan Zhao, Nora A. Foroud, Daqun Liu, Xiumei Yu.

**Funding acquisition:** Xiumei Yu.

**Project administration:** Xiumei Yu.

## Supporting information

**S1 Fig. Classification of FBK proteins in wheat based on different functional domains.** F-box, the protein with F-box domain; Kelch, F-box protein having Kelch domain; PAS, FBK protein with PAS domain that was named after three proteins that it occurs in: Per-period circadian protein, Arnt-Ah receptor nuclear translocator protein and Sim-single-minded protein; PAC, FBK protein with PAC domain that usually appears at the C-terminus of the PAS motif.

**S2 Fig. WebLogo generated by alignments of the F-box (A) or Kelch (B) domains of wheat FBKs.** The F-box or Kelch motifs were retrieved from 74 wheat F-box proteins. The overall height of every stack is indicative of sequence conservation at the given position within the motif, whereas the height of the letters within each stack is indicative of the relative frequency of the corresponding amino acid. The bit score represents the information content for each position. Asterisks mark the conserved residues.

**S3 Fig. Chromosomal distribution of wheat FBK genes.** The chromosomes were drafted to proportion and the chromosome numbers were indicated at the top of each stave. Chromosomal distances were given in megabases (10 Mb). The gene names were listed at the right side of each chromosome corresponding to the position of each gene. Tandemly duplicated genes were shown in colored boxes. Segmental duplications were shown in coloured blocks.

S1 Table. Primer sequences.

S2 Table. Characteristics of wheat, Arabidopsis, rice, sorghum and maize FBK proteins.

S3 Table. FPKM values of wheat FBK genes.

S4 Table. Screening of the candidate proteins interacting with TaAFR.

S5 Table. Bioinformatics analysis of the candidate proteins.

S1 File. Sequences of FBK proteins in wheat, Arabidopsis, rice, sorghum and maize.

